# FUNCTIONAL NETWORK CONNECTIVITY (FNC)-BASED GENERATIVE ADVERSARIAL NETWORK (GAN) AND ITS APPLICATIONS IN CLASSIFICATION OF MENTAL DISORDERS

**DOI:** 10.1101/867168

**Authors:** Jianlong Zhao, Dongmei Zhi, Weizheng Yan, Vince D. Calhoun, Jing Sui

## Abstract

Functional network connectivity (FNC) obtained from resting-state functional magnetic resonance imaging (fMRI) data have been commonly used to study mental disorders in neuroimaging applications. Likewise, generative adversarial networks (GANs) have performed well in multiple classification benchmark tasks. However, the application of GANs to fMRI is relatively rare. In this work, we proposed an FNC-based GAN for classifying brain disorders from healthy controls (HCs), in which FNC matrices were calculated by correlation of time courses derived from non-artefactual fMRI independent components (ICs). The proposed GAN model consisted of one discriminator (real FNCs) and one generator (fake FNCs), each has four fully-connected layers, and feature matching was implemented between each other to improve classification performance. An average accuracy of 70.1% with 10-fold cross-validation was achieved for classifying 269 major depressive disorder (MDD) patients from 286 HCs, at least 5.9% higher compared to other 6 popular classification approaches (54.5-64.2%). In another application to discriminating between 558 schizophrenia patients and 542 HCs from 7 sites, the proposed GAN model achieved 80.7% accuracy in leave-one-site-out prediction, outperforming support vector machine (SVM) and deep neural net (DNN) by 3-6%. To the best of our knowledge, this is the first attempt to apply GAN model based on fMRI data for mental disorder classification. Such a framework promises wide utility and great potential in neuroimaging biomarker identification.

## 1. INTRODUCTION

Functional connectivity (FC) has been commonly utilized to study mental disorders, which can reflect the organization and interrelationship of spatially separated brain regions. FC is widely applied in neuroimaging to identify potential biomarkers for predicting or classifying mental disorders. Mental disorders cause high socioeconomic burdens and many disease exhibit comorbidity between each other (1). However, diagnosis of mental disorders mainly depends on symptom scores from clinical interview, such as the Hamilton depressive rating scale (HDRS) and the positive and negative syndrome scale (PANSS), which lack reliable and objective biomarkers. There are no existing gold standards that can be used for definitive validation. FC has shown great potential to differentiate mental disorders such as schizophrenia and major depressive disorder (2, 3).

Many machine learning methods based on FC have been applied in the classification of mental disorder. For instance, support vector machine (SVM), linear discriminant analysis (LDA), and nearest neighbors (NN) methods have been successfully applied to discriminate mental disorders (2). However, traditional machine learning algorithms has been criticized for its poor performance on raw data because it requires expert experience to apply feature selection to acquire less redundant and more informative features. Recently, deep learning has achieved remarkable results in many research field (4-6). Deep learning is able to automatically learns from the pattern of data without feature selection and the accuracy of classification using deep learning on neuroimaging data is higher than traditional machine learning algorithms. For instance, (7) adopted stacked autoencoder to initialize its own weight based on pre-training to increase schizophrenia classification accuracy. (8) investigated the discriminant autoencoder network for multi-site classification of schizophrenia with fMRI. Therefore, the deep learning methods have demonstrated powerful diagnostic ability for mental disorder classification and provide better analysis of pathophysiology.

In particular, generative adversarial networks (GANs) have drawn increasing attention due to their capability to perform data generation and have been widely used in many fields, including image synthesis, reconstruction, segmentation, and classification(9). GANs have been recently proven successful on standard classification benchmark tasks (10, 11). The generated samples promote the discriminator’s classification ability through adversarial learning, especially in the case of small samples size. For instance, (11) proposed multiclass spatial–spectral GAN (MSGAN), the discriminator enhances classification performance by extracting the joint features of spatial and spectral information generated by two generators. The features for these problems are high-dimensional but the small sample size problem in medical imaging makes it challenging for the classifiers to learn a good decision boundary.

Inspired by this, in this work, we proposed an FNC-based GAN for classifying brain disorders from healthy controls (HCs), in which FNC matrices were calculated by correlation of time courses derived from non-artefactual fMRI independent components (ICs). Functional network connectivity (FNC) is able to reflect functional interactions among structurally segregated brain regions, known as brain networks (12). While FC is widely computed based on the predefined regions of interest (ROI), FNC estimated by group independent component analysis (group-ICA) demonstrates more reliable and sensitive in biomarker detection for psychosis (13).

To the best of our knowledge, this is the first attempt to apply GAN based on FNC to discriminate MDD vs HC (555 subjects, 269 MDD patients and 286 HCs from 4 sites) and SZ vs HC (1100 subjects, 558 SZ patients and 542 HCs from 7 sites). The proposed GAN model shows a GAN-based end-to-end classification and it combines FNC generation sample and classification into a unified optimization framework. In addition, the proposed GAN model defines adversarial objections between the generators and discriminator and uses adversarial learning to further improve classification performance of the discriminator. Our model alleviates the small size problem of FNC images by making full use of generated FNC samples and adversarial learning. Results showed that the GAN model achieved an average 70.1% accuracy with 10-fold cross-validation in MDD vs HC, at least 5.98% higher than five conventional methods and deep neural net (DNN) (54.5-64.2%). To validate the effectiveness of GAN model, we further applied it to a large-scale multi-site schizophrenia (SZ) dataset including 558 patients and 542 HCs from seven sites, achieving 80.73% accuracy in leave-one-site prediction, outperforming SVM and DNN by 3-6%, further demonstrating efficacy of the GAN approach.

## 2. MATERIALS AND METHODS

### 2.1. Participants

In this study, we used the 555 Chinese Han samples including 269 MDD patients and 286 HCs and these samples were derived from 4 sites, including the Henan Mental Hospital of Xinxiang(Site 1), the West China Hospital of Sichuan(Site 2), the Anding Hospital of Beijing(Site 3), and the First Affiliated Hospital of Zhejiang(Site 4). No significant group difference between HC and MDD was obtained in age or gender (age: p = 0.22; gender: p = 0.18). The DSM-IV based on SCID-P interviews was used to diagnose patients. HCs were interviewed using SCID-I/NP, and the first-degree relatives with any mental illness were excluded.

In this study, we also used 1100 Chinese subjects to test our method, including 542 HCs and 558 SZ patients from seven sites. Table 2 provides a more detailed statistical information.

**Table 1.** Demographic information of the HC/MDD database.

|  | MDD | HC | P-value |
| --- | --- | --- | --- |
| Number of scans | 269 | 286 | NA |
| Age (mean±std, yrs) | 32.8±10.6 | 31.7±10.4 | 0.22 |
| Gender (M/F) | 105/164 | 105/181 | 0.18 |

**Table 2.** Demographic information of the SZ/HC database.

|  | HC | SZ |
| --- | --- | --- |
| Number of scans | 542 | 558 |
| Age (mean±std, yrs) | 28.0±7.2 | 27.6±7.1 |
| Gender (M/F) | 276/266 | 292/266 |

### 2.2. Data Preprocessing and FNC Measure

The fMRI data for all subjects were preprocessed using SPM8 software (https://www.fil.ion.ucl.ac.uk/spm/), including the removal of the first 10 volumes, slice timing correction, motion correction, spatially normalized into standard MNI space, reslicing to 3 × 3 × 3 mm3 voxels, and spatially smoothing with a 6 mm full width half max (FWHM) Gaussian kernel.

For MDD patients and HCs, the fMRI data were decomposed into subject-specific spatial independent components (ICs) and its corresponding time courses using a spatially contrained ICA back-reconstruction approach called group ICA (group-ICA) implemented in the GIFT software (http://trendscenter.org/software/gift) which is robust to artifacts(18), resulting in 29 selected intrinsic connectivity networks (ICNs) from 100 group independent components, please see more details in (19). The time courses of selected ICNs were post-processed by detrending linear, quadratic and cubic trends, regressing out 6 realignment parameters and their temporal derivatives, despiking, and bandpass filtering between 0.01∼0.15 Hz using a 5th order Butterworth filter. FNC was computed as the pairwise correlation between any two ICN time courses for each subject, which was further used as input feature of the GAN model.

For SZ patients and HCs, the fMRI data were decomposed into subject-specific spatial independent components (ICs) and its time courses by performing group-ICA within the GIFT software(18), resulting in 50 selected intrinsic connectivity networks (ICNs) from 100 group independent components. The time courses (TCs) of selected ICNs are post-processed by detrending, regressing out head motion, despiking and lowpass filtering (<0.15 Hz) and the FNC matrices are calculated as the Pearson’s correlation in each pair of ICs, which was further used as input feature of the GAN model.

### 2.3. GAN Based on Feature Matching Classifier

As shown in Fig 1, a GAN architecture was applied for classification. Compared to unsupervised GAN model, the proposed GAN incorporated both labeled and generated data into the loss function that can be divided into three parts. The discriminator’s output layer has K + 1 classes, where K = 2 for the real class from data x and K + 1 class for the generated image. The loss function L is defined for each type of data (L_labeled_, L_unlabeled_, L_generated_), and the total loss is used to optimize the GAN model. The loss of L_unlabeled_ and L_generated_ made up the L_unsupervised_ loss that was trained without using the label information, while the GAN model used the label information to minimize L_supervised_ loss. All loss function is defined as below:

**Fig. 1.**
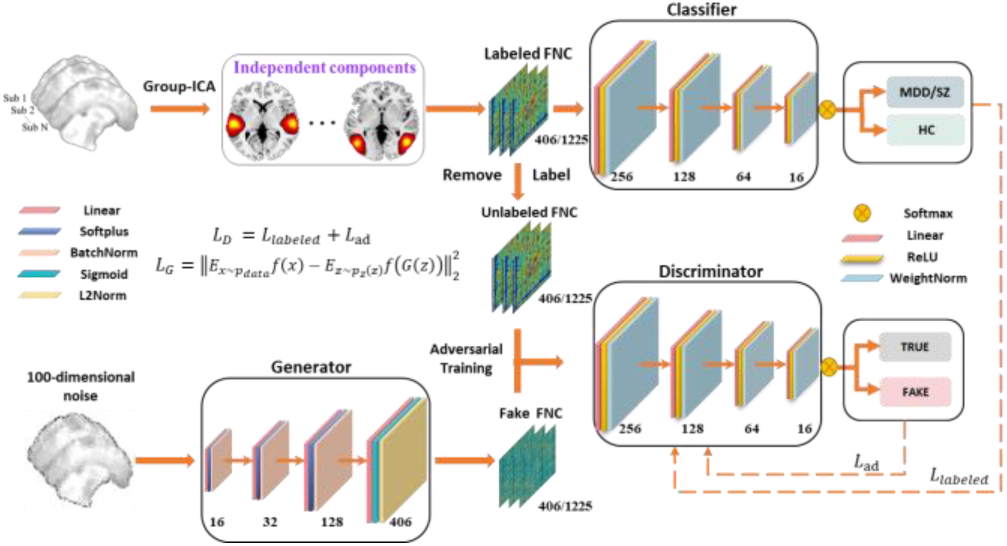
Overview of the GAN model for MDD vs HC and SZ vs HC classification. The GAN model was composed of a discriminator and a generator with four fully-connected layers.

**Fig. 2.**
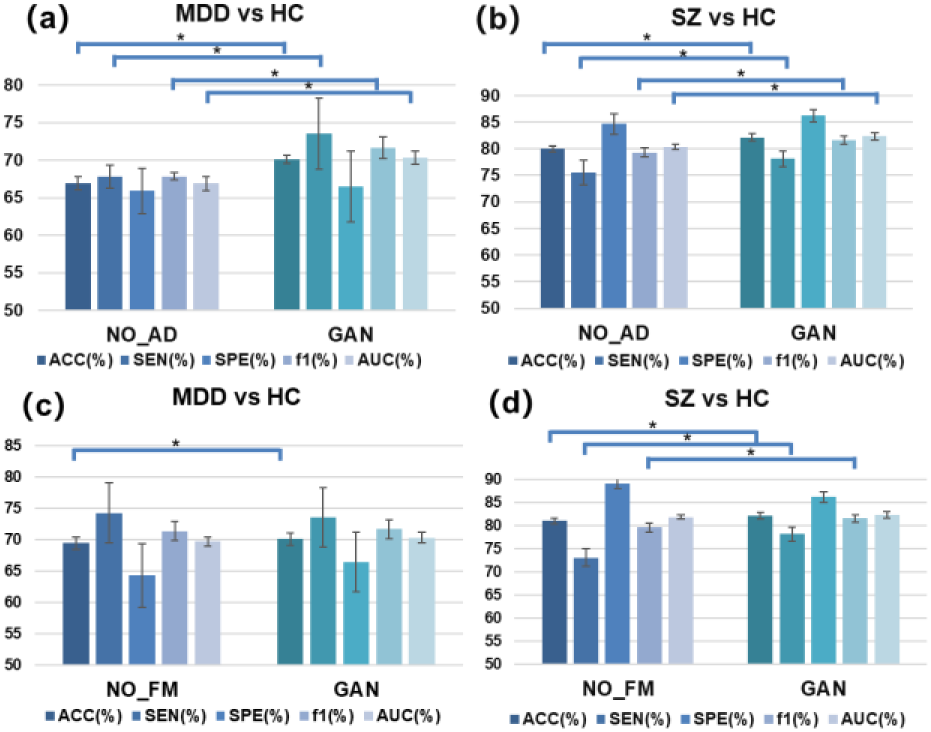
Effect of adversarial training and feature matching. No-AD: no adversarial training No-FM: no feature matching

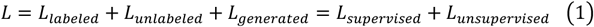

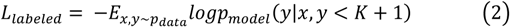

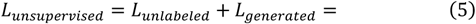

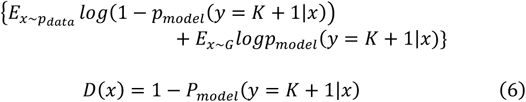

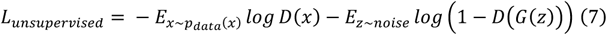

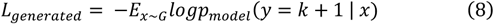

Where, x: input data; y: the label of the input data; P_data_: the distribution of the input data; G: the generated data; P_model_(. |.): the output probability of discriminator.

Feature matching can generate fake samples within the high-density region in feature space, which can split the bounds of different classes because of its continuity to further enhance classification performance (14). The generator was trained to match the feature value in intermediate layers of the discriminator.

Activations in an intermediate layer of the discriminator were denoted as f(x), our new objective for the generator can be defined as follows:

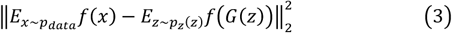

Previous studies revealed that the generated samples promote the discriminator’s classification ability through adversarial learning, especially in the case of small samples size (10, 11, 15) and feature matching is able to improve classification performance(10). Thus, we attempt to apply Feature Matching GAN based on FNC to discriminate MDD vs HC and SZ vs HC.

### 2.4. GAN model implementation

The GAN model was trained and evaluated by using Theano and Scikit-learn (https://scikit-learn.org/). The GAN model consisted of one discriminator and one generator that had four layers respectively. The output layer of the discriminator has K+1 classes, where K = 2 for the input pattern from HC and patient group, and the [K+1]th class is for generating fake images. The high-dimensional features of each subject may lead to overfitting on the training set. In order to reduce the overfitting susceptibility, L2 normalization and batch normalization were added to the generator to improve modal generalization. In addition, weight normalization was used to the output of each layer of the discriminator to prevent overfitting. The loss function of the GAN for the fine-tuning step is defined in 2.3.

The adam optimizer was adopted as minimizing the loss of the GAN model and a standard error back-propagation algorithm was used by training the GAN model with multiple layers. The batch size was set as 120 in the training process. In addition, to overcome overfitting, the weights were controlled with weight norm regularization. Different layers were attempted to the constructed architecture of the GAN model and results revealed that using four layers could obtain the optimal classification performance. The learning rate was set as 0.003. All trainings and experiments were completed on a standard workstation (Intel(R) Xeon(R) CPU E5-1650 v4 @ 3.60GHz, 6 CPU cores, 12GB NVIDIA GTX TITAN).

## 3. EXPERIMENT AND RESULTS

### 3.1. Ten-fold and leave-one-site-out Classification in MDD

To demonstrate the performance of the proposed method, we compared the proposed method with other state-of-art methods, including five conventional methods SVM, NN, Gaussian Process, Naive Bayes, and AdaBoost and a deep learning method DNN. 10-fold cross-validation strategy was used for estimating the generalization ability of the classifiers. In order to test the generalization of the model, we train the different models with the leave-one-site-out method. The classification results of the GAN model compared with the other six methods were summarized in Table 3 and Table 4. The result suggested that the proposed GAN method significantly outperformed other methods in terms of ACC, SEN, SPE, F1, and AUC measures. The t-distributed stochastic neighbor embedding (t-SNE) was used to visualize the GAN classification performance.

**Table. 3.** Performance of different methods in MDD/HC classification with 10-folds cross-validation

| Method | ACC(%) | SEN(%) | SPE(%) | f1(%) | AUC(%) |
| --- | --- | --- | --- | --- | --- |
| Nearest Neighbors | 54.54(0.92) | 55.85(0.92) | 53.13(0.94) | 56.08(0.87) | 54.48(0.93) |
| AdaBoost | 54.58(1.68) | 55.75(1.54) | 53.26(1.84) | 56.58(1.77) | 54.49(1.67) |
| Naive Bayes | 59.15(0.85) | 62.14(0.98) | 56.81(0.77) | 57.25(1.00) | 59.35(0.85) |
| Gaussian Process | 60.36(0.60) | 61.06(0.75) | 59.56(0.50) | 62.38(0.37) | 60.25(0.63) |
| Linear SVM | 62.81(0.71) | 61.90(0.59) | 64.19(0.92) | 66.73(0.66) | 62.51(0.71) |
| Deep Neural Net | 64.23(0.9) | 64.43(0.96) | 64.06(1.22) | 66.32(1.16) | 64.10(0.9) |
| **GAN** | **70.11(0.59)** | **73.53(4.73)** | **66.47(4.73)** | **71.65(1.47)** | **70.31(0.85)** |

**Table. 4.** Performance of leave-one-site-out method in MDD/HC classification

| Method | ACC(%) | SEN(%) | SPE(%) | f1(%) | AUC(%) |
| --- | --- | --- | --- | --- | --- |
| Nearest Neighbors | 55.50(1.96) | 57.25(2.48) | 53.85(4.64) | 55.50(4.38) | 55.55(1.19) |
| AdaBoost | 47.93(1.39) | 49.30(5.17) | 47.08(1.53) | 42.08(1.61) | 48.28(1.99) |
| Naive Bayes | 55.14(6.75) | 59.49(9.47) | 52.78(6.88) | 48.23(12.74) | 55.60(6.31) |
| Gaussian Process | 56.94(4.97) | 59.22(4.55) | 55.00(6.84) | 55.82(8.93) | 57.07(2.62) |
| Linear SVM | 55.86(6.2) | 58.95(2.67) | 53.68(8.99) | 52.43(15.8) | 56.13(4.76) |
| Deep Neural Net | 54.95(3.45) | 60.84(4.99) | 52.44(3.10) | 44.69(2.08) | 55.58(1.81) |
| **GAN** | **64.32(2.91)** | **61.46(3.05)** | **70.76(6.54)** | **70.45(1.54)** | **63.75(3.35)** |

### 3.2. The Classification of Ten-fold and leave-one-site-out in SZ

To validate the effectiveness of the GAN model, we evaluated the proposed GAN method on the task of SZ classification (542 HCs and 558 SZ). The proposed method was compared with six state-of-art methods using 10-fold cross-validation and leave-one-site-out method. Results were summarized in Table 5 and Table 6, demonstrating that the classification performance of GAN was significantly better than other methods. The proposed GAN model can be generalized to classify new site.

**Table. 5.** Performance of different methods in SZ/HC classification with 10-folds cross-validation

| Method | ACC(%) | SEN(%) | SPE(%) | f1(%) | AUC(%) |
| --- | --- | --- | --- | --- | --- |
| Nearest Neighbors | 64.6(0.64) | 69.1(0.80) | 61.59(0.58) | 61.03(0.8) | 64.75(0.64) |
| AdaBoost | 72.25(1.05) | 72.41(1.06) | 72.09(1.11) | 72.8(1.03) | 72.24(1.05) |
| Naive Bayes | 69.63(0.5) | 67.46(0.42) | 72.65(0.64) | 72.14(0.47) | 69.51(0.49) |
| Gaussian Process | 64.28(0.78) | 68.46(1.02) | 61.42(0.67) | 60.91(0.91) | 64.42(0.78) |
| Linear SVM | 77.08(0.9) | 77.37(1.04) | 76.79(0.88) | 77.43(0.86) | 77.08(0.9) |
| Deep Neural Net | 80.26(0.47) | 80.78(0.9) | 79.78(0.87) | 80.48(0.52) | 80.26(0.47) |
| **GAN** | **82.1(0.70)** | **78.14(1.51)** | **86.18(1.12)** | **81.57(0.82)** | **82.33(0.67)** |

**Table. 6.** Performance of leave-one-site-out methods in SZ/HC classification.

| Method | ACC(%) | SEN(%) | SPE(%) | f1(%) | AUC(%) |
| --- | --- | --- | --- | --- | --- |
| Nearest Neighbors | 64.27(5.93) | 67.67(8.22) | 61.77(9.80) | 61.66(5.98) | 64.39(5.15) |
| AdaBoost | 70.36(5.02) | 70.00(7.75) | 70.77(10.0) | 71.35(4.20) | 70.33(4.99) |
| Naive Bayes | 66.55(6.00) | 64.35(8.75) | 69.86(10.9) | 69.84(5.13) | 66.40(5.66) |
| Gaussian Process | 61.82(4.03) | 64.62(6.48) | 59.71(8.67) | 59.22(5.18) | 61.92(3.13) |
| Linear SVM | 74.91(2.86) | 74.14(7.89) | 75.78(8.62) | 75.83(1.92) | 74.87(3.05) |
| Deep Neural Net | 77.36(2.50) | 76.96(8.99) | 77.80(8.25) | 77.98(1.82) | 77.34(2.40) |
| **GAN** | **80.73(3.79)** | **81.8(3.74)** | **79.68(9.72)** | **80.76(3.91)** | **80.74(3.36)** |

### 3.3. Ablation experiment

Furthermore, adversarial training (AD) and FM in GAN could directly influence the learning capacity of the GAN method. Therefore, we had a comparison for the GAN models performance with adversarial training vs no adversarial training (No-AD) and feature matching vs no feature matching (No-FM). As shown in Fig. 5, the GAN model with adversarial training has increased performance from all evaluation criteria, compared to the GAN model with No-AD. The GAN model with FM has better classification performance compared to the GAN model with No-FM.

## 4. CONCLUSION

In summary, as far as we know, this is the first attempt to apply GAN based on FNC to discriminate MDD vs HC and SZ vs HC. We proposed a deep learning framework with GAN model, combined with adversarial training and feature matching, for brain disease diagnosis using the FNC, which can be directly used to classify new individual patients. Compared with 5 traditional methods and DNN approach, the GAN model achieved 5.9% higher in MDD classification and 1.8% higher in SZ classification, suggesting its utility as a potentially powerful tool to aid in discriminative diagnosis.

## Acknowledgment

This work was supported by the Natural Science Foundation of China(No. 61773380), the Strategic Priority Research Program of the Chinese Academy of Sciences (No.XDB32040100), Beijing Municipal Science and Technology Commission (Z181100001518005), the National Institute of Health (1R56MH117107, R01EB005846, R01MH094524, and P20GM103472) and the National Science Foundation (1539067).

